# Quantifying physiological trait variation with automated hyperspectral imaging in rice

**DOI:** 10.1101/2022.12.14.520506

**Authors:** To-Chia Ting, Augusto Souza, Rachel K. Imel, Carmela R. Guadagno, Chris Hoagland, Yang Yang, Diane R. Wang

**Affiliations:** Agronomy Department, Purdue University, West Lafayette, IN 47907 (USA); Institute for Plant Sciences, Purdue University, West Lafayette, IN 47907 (USA); Botany Department, University of Wyoming, Laramie, WY 82071 (USA)

**Keywords:** genetic diversity, growth traits, high-throughput phenotyping, hyperspectral imaging, nitrogen, *Oryza sativa*

## Abstract

Advancements in hyperspectral imaging (HSI) and establishment of dedicated plant phenotyping facilities have enabled researchers to gather large quantities of plant spectral images with the aim of inferring target phenotypes non-destructively. However, large volumes of data that result from HSI and corequisite specialized methods for analysis may prevent plant scientists from taking full advantage of these systems. Here, we explore estimation of physiological traits in 23 rice accessions using an automated HSI system. Under contrasting nitrogen conditions, HSI data are used to classify treatment groups with ≥ 83% accuracy by utilizing support vector machines. Out of the 14 physiological traits collected, leaf-level nitrogen content (N, %) and carbon to nitrogen ratio *(C:N)* could also be predicted from the hyperspectral imaging data with normalized root mean square error of predictions smaller than 14% (R^2^ of 0.88 for *N* and 0.75 for *C:N).* This study demonstrates the potential of using an automated HSI system to analyze genotypic variation for physiological traits in a diverse panel of rice; to help lower barriers of application of hyperspectral imaging in the greater plant science research community, analysis scripts used in this study are carefully documented and made publicly available.

**HIGHLIGHT:** Data from an automated hyperspectral imaging system are used to classify nitrogen treatment and predict leaf-level nitrogen content and carbon to nitrogen ratio during vegetative growth in rice.

## INTRODUCTION

Variation in plant traits reflect differences in genetics, the environment, and their interactions integrated over time (Allard and Bradshaw, 1964). Understanding these relationships could provide mechanistically-based insights into breeding climate-resilient crops for future climates. Conventional methods to measure plant traits, however, are largely destructive, time-consuming and often require technical skills specific to each particular method. Deriving trait data from imaging as a standardized means of non-destructive, high-throughput phenotyping has thus become of great interest to the plant research community over the past decade (Yang et al., 2014, 2020; Mir et al., 2019). Morphometric information, such as plant surface area and plant height, are standard data products that can be currently derived from Red-Green-Blue (RGB) images using established pipelines (Yang et al., 2014; Gehan et al., 2017; Berry et al., 2018; Kim et al., 2020). In contrast, pipelines for predicting physiological responses and/or biochemical traits from high-throughput imaging technologies have lagged (Pasala and Bb, 2020; Yang et al., 2020). Since these traits provide key insights to understanding responses of plants to abiotic factors, lowering the barriers to their systematic measurement across larger panels of genetic material is a valuable area of investigation (Silva-Perez et al., 2018).

Hyperspectral imaging (HSI) has shown promise as a potential high-throughput means of estimating physiological and biochemical traits in plants. HSI systems have a light source, objective lenses, an imaging spectrograph, hyperspectral sensor(s), and a computer. Results are stored as quantitative electrical signals derived from a vast number of images, each corresponding to the reflectance value – the ratio of reflected radiant flux to the incident flux – of any given wavelength, ranging between 400 to 2500 nm (Sarić et al., 2022). As early as the 1970s, plant scientists have documented the relationship between leaf traits (e.g., thickness, water content, presence of wax and hairs, and age) and hyperspectral reflectance (Gausman and Allen, 1973; Grant, 1987). For example, low reflectance in the visible light (VIS, 400 – 700nm) region is due to the absorption of light by photosynthetic pigments (Sarić et al., 2022). Reflectance in the near infrared (NIR, 700 – 1100nm) region is influenced by the arrangement of mesophyll tissues of leaves (Rouse et al., 1974), and the two troughs observed in the short-wave infrared (SWIR, 1000 – 2500nm) region are affected by water inside the plant cells (Cotrozzi et al., 2020). These findings support the utility of spectral features as surrogates for plant physiological and biochemical traits as part of high-throughput phenotyping systems.

Advancements in computing have facilitated exploration of HSI data in crops and the possible meaningful relationships with target plant traits. Early on, the properties of high dimensionality and multicollinearity precluded classical regression techniques for analyzing these large-scale data. Methods such as partial least squares regression (PLSR) are needed to quantify traits from HSI data while the recent use of machine learning (ML) algorithms to HSI-derived data has been shown to vastly improve predictions (Mir et al., 2019; Mishra et al., 2020; Arias et al., 2021). For example, support vector machines (SVM) have been recognized as one of the effective imaging classification algorithms (Noble, 2006; Gewali et al., 2019). Despite the capacity of PLSR and ML algorithms to handle large volumes of data, wavelength selection prior to building prediction models remains critical (Gewali et al., 2019; Yu et al., 2020). Principal component analysis (PCA) is one example of an unsupervised method to select wavelengths. By generating a set of principal components, with each PC a linear combination of the input variables, the dimensionality of HSI data is reduced (Gewali et al., 2019; Yu et al., 2020). Wavelength selection can also be achieved by supervised methods, such as the ReliefF algorithm. This is a ranking-based method that calculates importance score of each wavelength by considering the similarity and dissimilarity between wavelengths (Ren et al., 2020). Both PCA and ReliefF algorithms are wavelength selection methods that aim to enhance model performance while retaining the most important information from the original HSI data.

There are several examples of HSI products informing the quantitative and qualitative analysis of crop responses to abiotic factors in controlled-environment phenotyping facilities. The response of maize under drought has been studied in PHENOVISION in Ghent University, Belgium (Asaari et al., 2019; Mertens et al., 2021). At the phenotyping facility at University of Nebraska-Lincoln and The Plant Accelerator at University of Adelaide, researchers also found that HSI-derived data can be used to predict leaf water content in two varieties of maize and four varieties of wheat (Ge et al., 2016; Pandey et al., 2017; Bruning et al., 2019). Pandey *et al*. (2017) reported that macronutrients (nitrogen, phosphorous, potassium and sulfur) could be estimated from HSI data in maize (B73 inbred line) and soybean (cv. Thorne). Finally, Bruning *et al*. (2019) demonstrated that HSI data could be used to generate nitrogen distribution maps at the whole-plant level in four varieties of wheat, leveraging the fact that HSI can be spatially explicit.

Cultivated Asian rice (*Oryza sativa*), consumed by more than half of the population in the world, has also been investigated for its spectral features under field and controlled environment conditions (Muthayya et al., 2014; Arias et al., 2021). For example, Din *et al*. (2017) found that leaf area index (LAI) at vegetative stage of one variety from *japonica* subpopulation could be estimated from Vegetation Indices derived from hyperspectral data. These experiments were carried out in the field using a spectroradiometer to collect canopy-level data. Spectral data has also been used to detect common leaf diseases across four varieties of rice grown in greenhouse conditions (Feng et al., 2021). The leaves were first sampled and then placed in a HSI system. Yu *et al*. (2020) developed leaf-level nitrogen content models from spectral data on a single *japonica* variety and a single *indica* variety under field conditions. Their canopy-level HSI data was collected with a spectroradiometer, and Vegetation Indices were used as model input. These previous studies on rice focused on a small number of accessions, leaving the utility of high-throughput spectral data from diverse germplasm yet to be demonstrated.

Here, we evaluate the applicability of an HSI system to *O. sativa* using 23 genetically diverse accessions from two deeply diverged subpopulations under two levels of nitrogen application. Known variability in genetics and environment (i.e. treatments) is leveraged in the experimental design to drive the extent of potential phenotypic variation. Wavelength selection methods, such as PCA and ReliefF algorithm, are used prior to building SVMs for classification and PLSR for quantification, respectively. Our overall goal is to test the utility of data derived from the HSI system for predicting a suite of plant physiological traits that reflected diverse aspects of plant growth. Specific objectives of this work are to (1) assess whether HSI data can classify genotypic and treatment groupings and (2) understand the types of plant traits that have the most potential to be predicted using canopy-level HSI data.

## MATERIALS AND METHODS

### Plant materials

A set of 23 bio-geographically diverse accessions from two publicly-available, purified germplasm collections, the Rice Diversity Panel (RDP) 1 and RDP 2 (McCouch et al., 2016), were evaluated for this study. These 23 lines encompassed 15 *indica* and eight *tropical japonica* accessions that originated from 17 countries across Asia, West Africa and the Americas (To limit potential confounding effects of development on other traits of interest, accessions were selected out of the tropically-adapted and phenologically-similar RDP subset screened by Wang *et al*. (**Table S1** and **Figure S1**). 2016). Seeds were obtained from the USDA-ARS, Dale Bumpers National Rice Research Center, Stuttgart, Arkansas, Genetic Stocks Oryza Collection (www.ars.usda.gov/GSOR).

### Growth conditions

The selected diversity panel was raised at the Ag Alumni Phenotyping Facility (AAPF), a controlled-environment high-throughput plant phenotyping facility at Purdue University (West Lafayette, Indiana, U.S.A.) for 94 days during the Summer and Fall of 2020. The facility had a fully automated growth chamber (Conviron®, Winnipeg, Canada) and weight-based irrigation system (Bosman Van Zaal, Aalsmeer, The Netherlands). Three conveyer belts with 32 positions *per* belt were allocated to this study. Of the 96 total positions, two were designated as “purge pots,” *i.e.*, pots into which the system flushes solutions in between changing fertigation/irrigation regimes. The remaining 94 positions were occupied by 22 genotypes x 2 replicates x 2 nutrient levels and one final genotype (cv. Cybonnet) x 3 replicates x 2 nutrient levels (described below). The temperature setpoint in the chamber was 26/22 °C day/night, relative humidity at 60%, and photosynthetically active radiation (PAR) levels were recorded between 550-600 µmol photon *m^-2^ s^-1^*. The environment was additionally tracked by affixing Lascar EL-USB-2-LCD Data Loggers to seven randomly selected pots at the sowing time. They recorded temperature and relative humidity every 10 minutes (**Figure S2a**). Average temperature and humidity from the loggers across the experimental period were 29.48±0.22/23.34±0.09 °C day/night (mean±SE), relative humidity at 64.62±0.69% (mean±SE). Lighting conditions followed long day (14 h day/10 h night) scheduling: lights turned on daily at 0600 h and turned off at 2000 h. Two seeds were sown *per* pot (6L in volume) in horticultural substrate, which was made up of a mixture of Profile Porous Ceramic Greens Grade and Berger BM6 All Purpose at a one-to-one ratio by volume. Plants were hand-watered until 10 days after sowing (DAS), at which point the seedlings were thinned and a weight-based automated irrigation was initiated. The experiment was designed with two nutrient treatment levels: high (300 ppm nitrogen, N1) and low (50 ppm nitrogen, N2). The fertigation solution was created by mixing Peters Excel 15-5-15 Cal Mag Special in reverse osmosis (RO) water. Each morning prior to chamber lights turning on, plants were irrigated to a preset weight with RO water. This target weight was increased by about 1.15 times during the experiment to account for the increase in transpirational demand of the growing plants (**Figure S2b**). RO water irrigation occurred every day except on scheduled days when a fixed volume of fertigation solution (either high or low concentration, depending on the treatment) was applied instead of RO water. This RO water irrigation and fertigation regime was theoretically designed so that each plant should receive enough water to meet individual transpirational demands while also receiving the fertilizer amount prescribed by their treatment.

### Imaging and image processing

Plants were imaged approximately three times *per* week beginning on 20 DAS and continuing until the end of the experiment using the automated imaging booth in AAPF. During each imaging event, one side-view and one top-view images were acquired. The HSI cameras used a scanning range that encompassed the VIS to NIR region (VNIR; 400 - 1000 nm, MSV 500 VNIR Spectral Camera, Middleton, Spectral Vision, WI, USA) and the SWIR region (Specim, Oulu, Finland); thus, generation two hypercubes of data image. White and dark reference tests were conducted for post-processing of the rice HSI image, where the relative light reflectance was estimated for each wavelength. This step was based on the work by Zhang *et al*. (2019). For the white reference, two panels of known material - one for the VNIR and another for the SWIR cameras - and spectral signature was scanned for both side and top cameras with all lights inside the imaging chamber on. The dark reference test was conducted with lens shutter closed and no lights present. The relative light reflectance (%) for each plant was calculated based on the normalized difference between these tests, as seen in Equation 1.

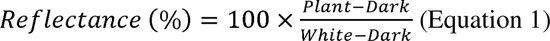

where,

Plant - the raw digital number measured by the HSI cameras for the rice plants

Dark - the raw digital number measured by the HSI cameras during the dark reference tests

White - the raw digital number measured by the HSI cameras during the white reference tests

For the VNIR hypercube, the rice plants were segmented out of the background using the typical attenuation between Red Edge wavelengths reflectance. The SWIR hypercube segmentation was done using the SURF image registration (Bay et al., 2008) to map the rice plants from the segmented VNIR hypercube (fixed image) using the VNIR-segmented rice plant as the fixed image. After the segmentation, the average light reflectance and fifty Vegetation Indices for each plant and scan were calculated and stored in spreadsheet files. HSI were processed by using a proprietary hyperspectral image processing script in MATLAB (MATLAB, 2018).

### Growth and physiology measurements

Pre-dawn gas exchange measurements were taken in the facility growth chamber during early vegetative growth between 0430-0530 h prior to chamber lights turning on using LI-6800 Portable Photosynthesis System (LI-COR, Lincoln, NE, USA) (**Table 1**). Leaf-level photosynthesis *(A, µmol m^-2^s^-l^*^·^), stomatal conductance to water vapor *(gsw, mol m^-2^s^-l^*), and nighttime transpiration *(E, mol m^-2^s^-l^*) were extracted from the measurements. Environmental conditions in the cuvette matched ambient conditions in the growth chamber: reference CO_2_, 415 µmol mol^-1^; vapor pressure deficit, <1.5 kPa (average VPD = 1.09 kPa); PAR, 0 µmol photon *m^-2^s^-l^*; and leaf temperature was 26.87±0.016 °C (mean ± SE). As rice leaves are generally too narrow to cover the entire cuvette, leaf width was first measured with a digital caliper prior to each measurement in order to adjust gas exchange measurements to the observed leaf area. A/C_i_ curves were collected in the facility growth chamber using both LI-6800 and LI-6400XT Portable Photosynthesis Systems (LI-COR, Lincoln, NE, USA) during 59-63 DAS and 87-91 DAS (mid-tillering and late vegetative stages, respectively) with constant PAR of 1200 µmol m^-2^ s^-1^ and reference CO_2_ concentrations were set in the following order: 415, 300, 200, 100, 50, 10, 415, 415, 600, 800, 100, 1200, 415 µmol mol^-1^. Light response curves were initially conducted on randomly selected plants to determine the PAR level for running A/C_i_ curves. A/C_i_ curves were collected between 1000 and 1400 h on the youngest fully expanded leaf as indicated by the emergence of the leaf collar. Leaf temperatures were between 25 and 27 °C and relative humidity was maintained between 50-70%. After A/C_i_ curves were run during 59-63 DAS, the leaf used for each curve was excised and its area measured using a leaf area meter (LAI-2200C; LI-COR, Lincoln, NE, USA). Leaves were oven dried for at least five days at 65 °C and their dry weights recorded. They were then finely ground using a mortar and pestle and subsequently analyzed for carbon *(C,* %) and nitrogen *(N,* %) content using the FlashEA® 1112 Nitrogen and Carbon Analyzer for Soils, Sediments and Filters (Thermo Scientific, CE Elantech, Lakewood, NJ) with two replicates *per* leaf (30 mg *per* replicate). The equipment was operated according to the flash dynamic combustion method, and resultant signals were translated into the percentage of carbon and nitrogen by the Eager 300 software. During 67-91 DAS, chlorophyll *a* fluorescence was measured at predawn (0430-0530 h) and midday (1000-1400 h) conditions with a hand-held fluorometer (Fluorpen FP110, Photon System Instruments, Drásov, Czech Republic) on the youngest, fully expanded leaf *per* plant on three different areas of the leaf blade to minimize possible variation in the efficiency of PSII due to the spatial response to sudden environmental changes in monocotyledons (Oberhuber et al., 1993): the basal one-third, the middle one-third, and the distal one-third. Measurements of Fv/Fm, the maximum efficiency of PSII in dark acclimated leaves, were taken according to Murchie and Lawson (2013). The measuring light of the FluorPen was set at approximately 1500 µmol photons m^-2^ s^-1^ throughout the experiment. Then, we applied a saturation pulse at approximately 2200 µmol photons m^-2^ s^-1^ to obtain Fv/Fm *(pQY),* whereas those taken at midday were the maximum efficiency of PSII in light conditions, *Fv’/Fm’(mQY)* (Henriques, 2009). Total above-ground biomass from all plants were harvested at the end of the experiment on 94 DAS, oven-dried at 65 °C for at least five days, and their dry weights recorded. **Table S2** summarizes the relationship between the timing for physiological trait measurements and the imaging dates.

**Table 1.** Overview of growth and physiology measurements.

| Abbreviation in text | Abbreviation in figure | Units | Description |
| --- | --- | --- | --- |
| A | A | $\mu\text{mol m}^{-2} \text{ s}^{-1}$ | pre-dawn leaf-level net assimilation |
| C | C | % | carbon |
| C:N | CN | - | leaf-level carbon to nitrogen ratio |
| **E** | E | $\text{mol m}^{-2} \text{ s}^{-1}$ | pre-dawn leaf-level transpiration |
| **FB** | FB | g | final biomass |
| **gsw** | gsw | $\text{mol m}^{-2} \text{ s}^{-1}$ | pre-dawn leaf-level stomatal conductance to water |
| **J<sub>max</sub>** | J<sub>max</sub> | $\mu\text{mol m}^{-2} \text{ s}^{-1}$ | light-saturated potential rate of electron transport |
| **log(E)** | log(E) | - | log transformed pre-dawn leaf-level transpiration |
| **log(gsw)** | log(gsw) | - | log transformed pre-dawn leaf-level stomatal conductance to water |
| **mQY** | mQY | - | midday quantum yield |
| **N** | N | % | nitrogen |
| **pQY** | pQY | - | pre-dawn quantum yield |
| **SLA** | SLA | $\text{cm}^2 \text{ g}^{-1} \text{ dw}$ | specific leaf area with respect to dry weight |
| **SLA_C** | SLA_C | $\text{cm}^2 \text{ mg}^{-1} \text{ C}$ | specific leaf area with respect to carbon |
| **SLN** | SLN | $\text{mg N cm}^{-2}$ | specific leaf nitrogen |
| **V<sub>cmax</sub>** | V<sub>cmax</sub> | $\mu\text{mol m}^{-2} \text{ s}^{-1}$ | the maximum carboxylation rate limited by Rubisco |

### Data analysis

Data were formatted and analyzed in R 4.1.1 (R Core Team, 2021) with packages dplyr (Wickham et al., 2021) and reshape2 (Wickham, 2007). Plots were made with package ggplot2 (Wickham, 2016) or in base R environment. The code for each physiological trait model can be accessed through GitHub (https://github.com/To-Chia/rice_imaging_ms).

#### Physiological trait collection

From the physiological trait measurements, we derived specific leaf area *(SLA,* cm^2^g^-1^), CN ratio *(C:N),* specific leaf area with respect to carbon *(SLAC* (cm^2^ mg^-1^ (C)) and specific leaf nitrogen *(SLN,* mg (N) cm^-2^). The summary statistics are in **Table S3**. Histograms and normal density distributions estimated from the data were plotted to decide whether the data needed to be transformed. *E* and *gsw* had right-skewed distributions (**Figure S3** and **Figure S4**) and thus were log-transformed (denoted as *log(E)* and *log(gsw),* respectively), prior to conducting the correlation tests and a PCA. Pearson’s correlation coefficient was calculated for all pairwise combinations of the physiological traits and plotted with package corrplot (Wei and Simko, 2021) (**Figure S5a**). Data for *SLA, C* and *N* on six rice plants were unavailable and were mean-imputed before the pairwise correlation was calculated. The same methods were applied to *C:N, SLA_C*, and *SLN*. A/C_i_ curves were first assessed for quality control (QC) using the PEcAn.photosynthesis package (https://pecanproject.github.io/modules/photosynthesis/docs/articles/ResponseCurves.html). For the 89 curves that passed QC, V_cmax_ and J_max_ were estimated using the R plantecophys package (Duursma, 2021). PCA based on correlation matrix was conducted on accession-mean data using the *prcomp* function in base R (R Core Team, 2021). For unavailable data, their replicates were used to represent the accession-mean value. Agglomerative hierarchical clustering was performed on the Euclidean distance matrix from accession-mean data. Clustering was based on the average linkage method and plotted with package dendextend (Galili, 2015).

#### Hyperspectral imaging data

Five Vegetation Indices heatmaps related to chlorophyll content and water availability were calculated from processed reflectance data (**Figure 1**). These included the Water Band Index (WBI) (Peñuelas et al., 1993), Short-wave infrared Normalized Difference Vegetation Index (swirNDVI) (Mayer and Scribner, 2002), Normalized Difference Water Index (NDWI) (Gao, 1996), Normalized Difference Vegetation Index (NDVI) (Roberts et al., 2004) and Moisture Stress Index (MSI) (Rock et al., 1986).

**Figure 1.**
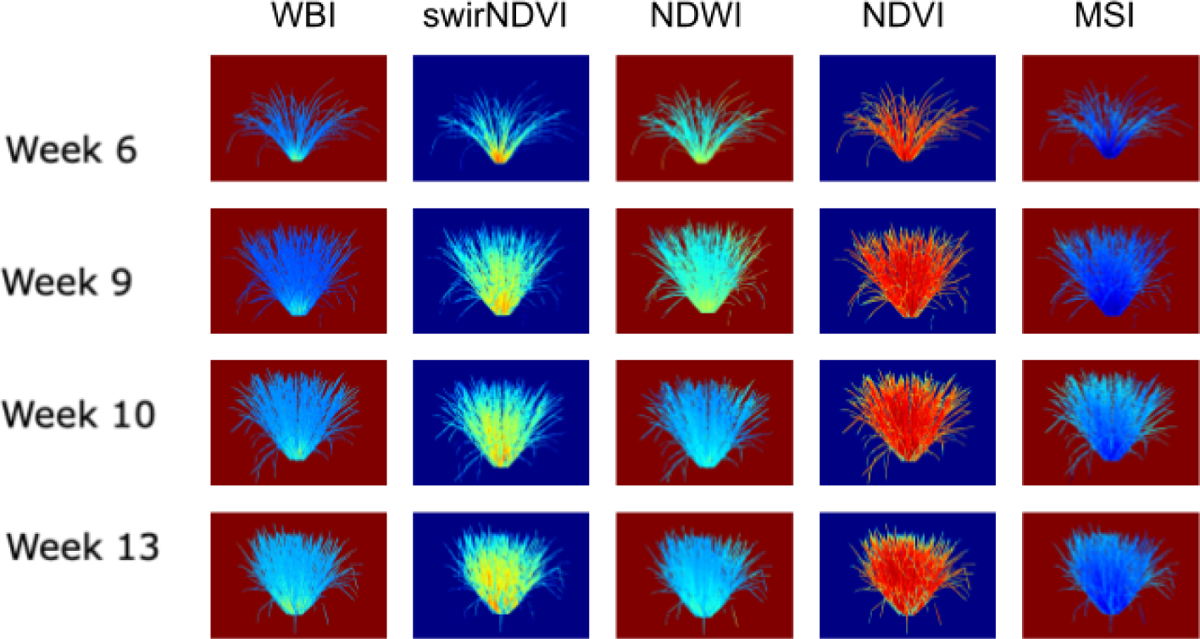
Example of hyperspectral images used for analyses. Side-view images of an individual replicate of the rice accession, Peh-Kuh-Tsao-Tu (NSFTV130 from Taiwan). From top to bottom, the images are taken from Weeks 6, 9, 10 and 13. From left to right, the images were colored to indicate the following Vegetation Indices: Water Band Index (WBI), short-wave infrared Normalized Difference Vegetation Index (swirNDVI), Normalized Difference Water Index (NDWI), Normalized Difference Vegetation Index (NDVI) and Moisture Stress Index (MSI).

To have stable signals for modeling, imaging events that occurred within the same weeks were averaged (**Table S2**). The averaged datasets were referred to as Week 6, 9, 10 and 13. PCA with variance-covariance matrix was conducted on the averaged HSI data. From the results of PCA, wavelengths that had the top ten loadings of the PCs that cumulatively accounted for >90% of total variance were selected and termed as Wsvm (**Table S4**).

#### Signal variation in HSI data through time

Based on the results from PCA, SVMs were trained to quantify the prediction accuracy of treatment groupings from W_SVM_ using the package e1071 (Meyer et al., 2021). First, the classifier of the training week was built with W_SVM_. Then, predictions made on the evaluation week were achieved by selecting W_SVM_ in the training week dataset from the evaluation week. We utilized radial basis function (RBF) kernels and linear kernels. Grid-search of the parameters for both kernels were conducted with 23-fold cross validation. The ranges of the parameters were referenced from Hsu *et al*. (2016) and both rough and fine grid-searches were conducted. In the rough grid-search of RBF kernels, parameters *cost* and *gamma* were searched within the range of 2^-5^ and 2^15^, 2^-15^ and 2^3^, respectively, both with an interval of 2^2^. The ranges of parameters in the fine grid-search were the best parameters in the rough grid-search minus and plus 2^0.5^, with an interval of 2^0.1^. When the best parameters in the rough grid-search fell on the border, the ranges of the parameters for the fine grid-search were conducted using the value plus or minus 2 within the specified ranges. Parameters found in the fine grid-search were adopted only if the model accuracy was higher in the fine grid-search than the rough grid-search; in some cases, the accuracy was the same. The parameter, *cost,* which was the only parameter in a linear kernel, was determined with the same method as *cost* in the RBF kernels.

#### Prediction of physiological traits with HSI data

To select feature wavelengths, termed as W_PLSR_, a regression based ReliefF algorithm in package CORElearn (Robnik-Sikonja and Savicky, 2021) was leveraged. The algorithm took conditional dependencies into account (Robnik-Šikonja and Kononenko, 2003) and was therefore an appropriate method to calculate the attributes values of the wavelengths for each physiological trait. The default estimator, “ReliefFexpRank,” was used and the number of iterations was the number of observations multiplied by 50. W_PLSR_ was composed of wavelengths with the top 300 attributes values and they were the input of a single­response (*i.e*., a physiological trait) PLSR models. The R package, pls (Liland et al., 2021), was used and the protocol for building the models was adapted from Burnett *et al*. (2021).

Specifically, models were fit using the classical orthogonal scores algorithm. Eighty percent of the full dataset was treated as the calibration dataset and the remaining was the validation dataset. Sampling was done with the criterion that each treatment level contributed equally to the datasets (*i.e.*, the full-data was grouped by treatments prior to sampling). Calibration datasets were used to determine the number of components in the final models by selecting the lowest root mean square error of prediction (RMSEP) in leave-one-out cross-validation. Final models were used to predict trait values using HSI data from validation sets. We calculated coefficient of determination *(R^2^,* Equation 2), root mean square of error in predictions (RMSEP, Equation 3) and normalized root mean square of error in predictions (%RMSEP, Equation 4) as performance metrics.

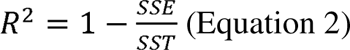

where *SST* is the sum of squares of the response and *SSE* is the sum of squares errors of the cross-validated predictions in the calibration dataset *(i.e,* predicted residual error sum of squares, PRESS) or predictions in the validation dataset.

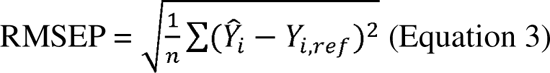

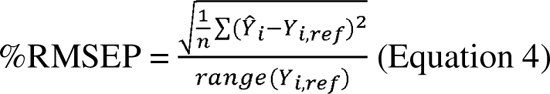

where *n* was the sample size, Ŷ_i_ was the predicted value estimated from the cross-validation in the calibration dataset or the predicted value in the validation dataset. *Ŷ_i,ref_* was the measured value. To estimate overall model performance, 95% confidence intervals derived from jackknife permutation during model calibration were used.

## RESULTS

### Treatment and subpopulation effects on multivariate physiological traits

Examining pairwise relationships among the traits, we found that all traits had significant correlations (alpha = 0.01) with at least one other trait, except for pre-dawn leaf-level net assimilation (*A*, µmol m^-2^ s^-1^) (**Figure S5b**). As expected, traits that reflected similar aspects of plant physiology were more correlated. For example, nitrogen content (*N*, %) and leaf-level carbon to nitrogen ratio *(C:N)* had a strong negative correlation (−0.96), log transformed pre-dawn leaf-level transpiration *(log(E))* and log transformed pre-dawn leaf-level stomatal conductance to water *(log(gsw))* were perfectly positively correlated, and *pQY* (Fv/Fm) and *mQY* (Fv’/Fm’) had a weak positive correlation (0.48). We found that overall health status of the rice plants improved with nitrogen enrichment, as reflected by higher *pQY* and *mQY* in N1 (high nitrogen) than in N2 (low nitrogen) treatments. *N* was also found to have moderate and weak positive correlations with *pQY* and *mQY,* respectively (0.53 and 0.41). Since *N* and *C:N* had a negative correlation, *C:N* had moderate and weak negative correlations with *pQY* and *mQY,* respectively (−0.55 and −0.44). While we did not detect a direct relationship between carbon (*C*, %) and *N*, we found specific leaf nitrogen *(SLN,* mg N cm^-2^) had a moderate negative correlation (−0.72) with specific leaf area with respect to carbon *(SLA C,* cm^2^ mg^-1^C). It was worthy to note that specific leaf area with respect to dry weight *(SLA,* cm^2^ g^-1^ dw) had correlations with most of these traits; it had a weak negative correlation with *C:N, SLN,* and *log(E)* (−0.46, −0.34, and −0.27) and a weak positive correlation with *N*, *SLA_C*, *mQY* and *pQY* (0.49, 0.42, 0.41, and 0.29). On the other hand, final biomass (*FB*, g) only had a weak negative correlation with *log(E)* and *log(gsw)* (−0.35 for both), suggesting that dry matter accumulation of rice in this experiment may have been more associated with water status than with nutrient status.

Due to the correlated nature of the physiological trait data described above, we employed several multivariate approaches to evaluate potential subpopulation and treatment effects on high-level data structure. PCA on accession-mean trait data suggested both treatment and subpopulation effects with PCs 1 and 2, which together accounted for 52% of total variance (**Figure 2a**). PC1 and PC2 reflected treatment and subpopulation effects, respectively. The traits, *N* and *C:N*, contributed to PC1 more than other traits whereas *SLA_C* and *SLN* had more influence on PC2 than the other traits. However, we only observed a clear separation by treatment using hierarchical clustering (**Figure 2b**). Interestingly, the dendrogram revealed that N2 *indica* observations were more closely clustered with the N1 group (both *indica* and *tropical japonica*) than with the cluster that primarily contained N2 *tropical japonica,* suggesting that *indica* may be less responsive to nitrogen enrichment than *tropical japonica*.

**Figure 2.**
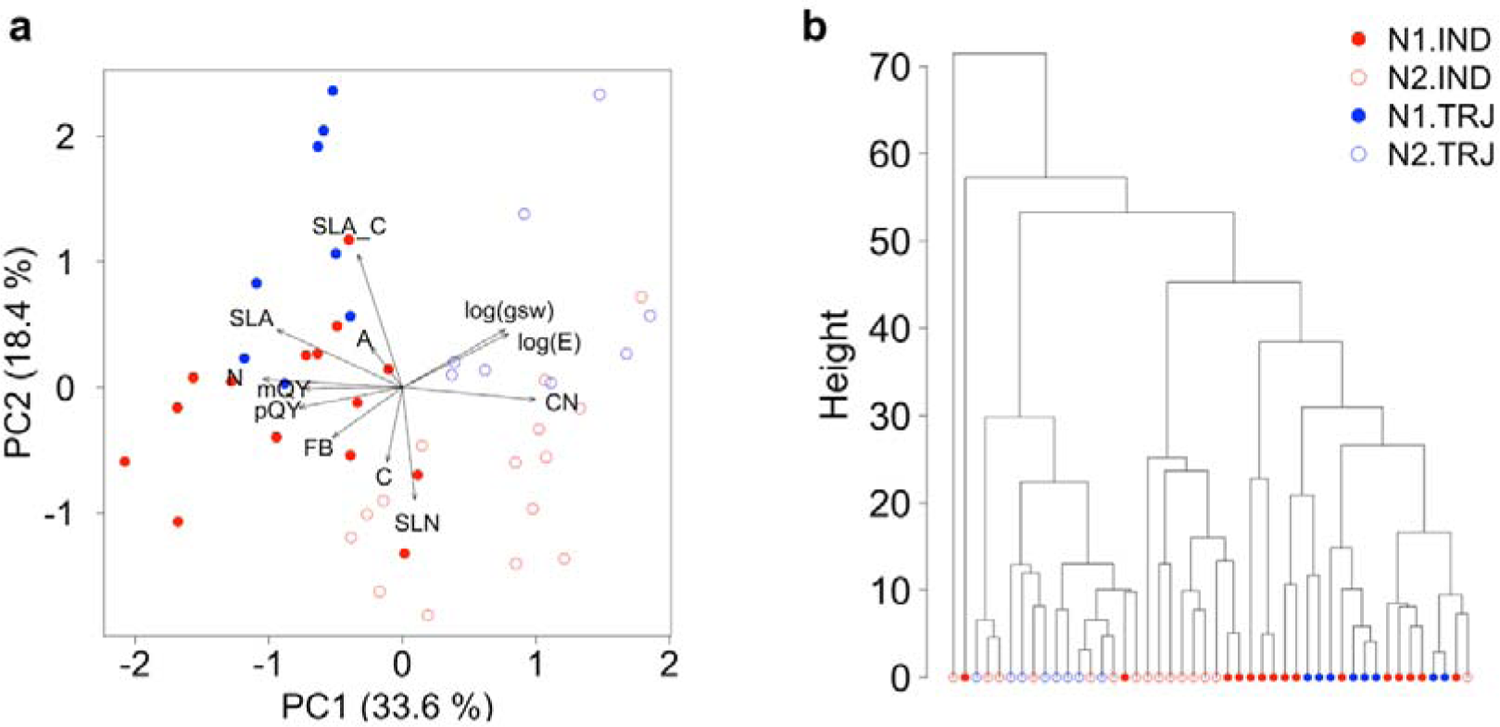
Treatment and subpopulation effects on physiological traits. (a) Principal components analysis biplot of PC1 (x-axis) and PC2 (y-axis). Blue and red colors indicate *tropical japonica* (TRJ) and *indica* (IND) subpopulations, respectively. Solid and empty circles represent high (N1) and low (N2) nitrogen treatments, respectively. SLA, C, N, CN, SLA_C, SLN, FB, mQY, pQY, A, log(E), and log(gsw) represent specific leaf area (cm^2^ g^-1^ dw), carbon (%), nitrogen (%), CN ratio, specific leaf area with respect to carbon (cm^2^ mg^-1^C), specific leaf nitrogen (mg N cm^-2^), final biomass (g), Fv’/Fm’, Fv/Fm, pre-dawn leaf-level net assimilation (μmol m^-2^s^-1^), log transformed pre-dawn leaf-level transpiration, and log transformed pre-dawn leaf-level stomatal conductance to water, respectively. (b) Agglomerative hierarchical clustering performed on accession mean physiological trait data.

### Variation in hyperspectral Vegetation Indices

Examining Vegetation Indices, we observed that WBI generally remained constant throughout the experimental period under both treatments and in both subpopulations (**Table S5**). MSI showed decreased trends for the two subpopulations under two treatments through time while NDWI revealed increasing trends. Taken together, these results might lead us to speculate whether plants suffered from water stress at the beginning of the experiment. However, the increase of NDVI from Week 6 to Week 9 suggested an increase of leaf area. Therefore, the observed trends of MSI and NDWI were most likely influenced by the occlusion of the rice leaves from each other. The results indicated that we need multiple Vegetation Indices to provide a comprehensive understanding of variation in traits.

### Effects of nitrogen on hyperspectral reflectance in rice

Due to limited insights from the Vegetation Indices, we next investigated whether HSI data could detect similar classes of subpopulation and treatment. Spectra indicated that the VIS region was largely absorbed by chlorophyll and other pigments, as would be expected for plants, with a small peak at the green portion of VIS light (520 - 600 nm), indicating that green light was not absorbed as strongly as light of other colors (*i.e.*, blue & red) by chlorophyll (**Figure 3**). Reflectance increased dramatically in 680 - 750 nm, a region known as the Red Edge. Interestingly, medians of reflectance values between 800 – 1300 nm increased consistently as rice grew from Week 6 to Week 13. This region was within the NIR region plateau (750 – 1300 nm), which is known to reflect changes in cell structure (Rouse et al., 1974), plant biomass and LAI in the whole plant scale (Din et al., 2017). In the SWIR regions of 1350 - 1500 and 1900 - 2100 nm, reflectance decreased, due to absorption of light by water inside the plants (Cotrozzi et al., 2020), forming two obvious V-shapes in the wavelength profiles. Finally, from 2200 - 2500 nm, the medians of reflectance values on Weeks 6 and 9 were higher than those on Weeks 10 and 13, which may result from large variation among individual plant spectra variation in this region (**Figure S6**).

**Figure 3.**
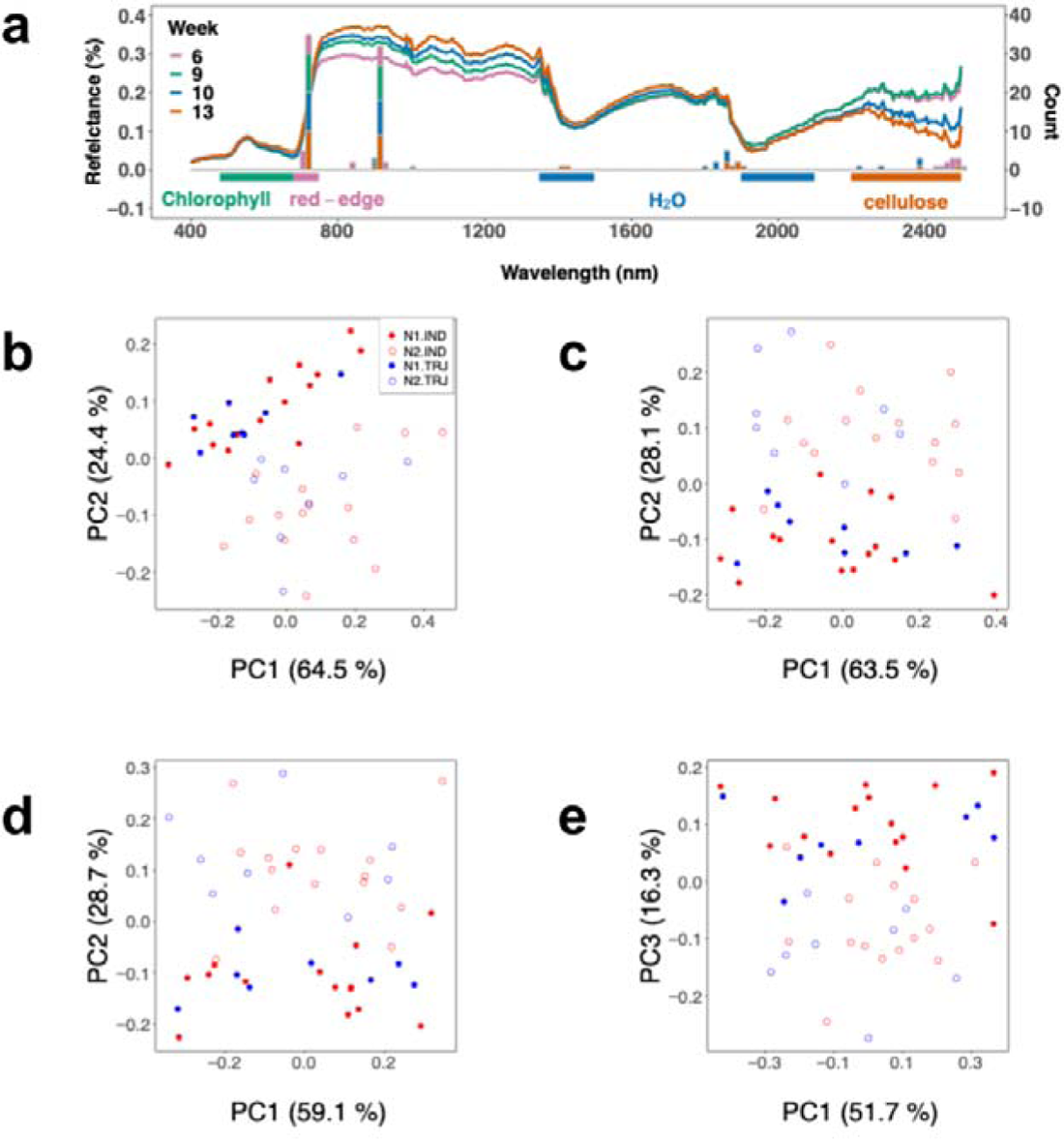
Variation in hyperspectral imaging (HSI) reflectance data over developmental time. (a) The medians of the accession-mean HSI reflectance data across weeks and the stacked bar plot of the key wavelengths identified from each week. Labelled colored horizontal bars represent wavelength regions with known associations with plant traits. Plots shown in (b), (c), (d), and (e) are principal component analysis conducted on accession-mean HSI reflectance data on Week 6, 9, 10 and 13, respectively.

PCA was applied on accession-mean HSI datasets to examine the potential effects of treatment and subpopulation (**Figure 3b-e**). Treatment effect was clear throughout the time period analyzed; separation of treatment levels was primarily determined by PC2 from Weeks 6 to 10 and by PC3 on Week 13. However, nitrogen treatment signals appeared to be lower on Week 13 than on earlier weeks; the variance explained by PC3 on Week 13 (16.3%) was lower than the variances explained by PC2 on Weeks 6 to 10 (24.4%, 28.1%, and 28.7% on Weeks 6, 9 and 10, respectively). Leveraging these PCA results, wavelengths were selected for use in support vector machines (SVM); these are referred to hereafter as W_SVM_ (see Methods for details). W_SVM_ detected in PC2 from Weeks 6 to 10 were similar to those detected in PC3 on Week 13 and were all centered around 715 nm (**Figure 3a** and **Table S4**) in the Red Edge region. This indicated that the treatment signals could be attributed to similar spectral regions across the full experimental period. A detailed examination showed that wavelengths around 715 and 910 nm accounted for the highest and the second highest proportions of W_SVM_, respectively. Wavelengths around 2200 – 2400 nm, the highly variable region, was found to contribute to the set of W_SVM_ as well. This was observed for all weeks except Week 9. Lastly, W_SVM_ of Weeks 10 and 13 included wavelengths around 1400 and 1800 nm, a region known to be informative of water absorption.

We next quantified treatment classification accuracy of these selected wavelengths by applying SVM with radial basis function (RBF) kernels and linear kernels. Results showed that W_SVM_ can classify nitrogen treatments for any one week using information from any other weeks (**Figure 4**). Overall, accuracies ranged from 0.63 to 0.91. We found that classification accuracy was greatest for Weeks 6 through 10 and poorest for Week 13. Accuracy of prediction made by Week 13 dropped as evaluation weeks became more temporally distant from the training week. The lowest accuracy using the Week 13 classifier was for Week 6 (0.7 for both kernels) while for Weeks 9 and 10, the accuracy was about 0.78.

**Figure 4.**
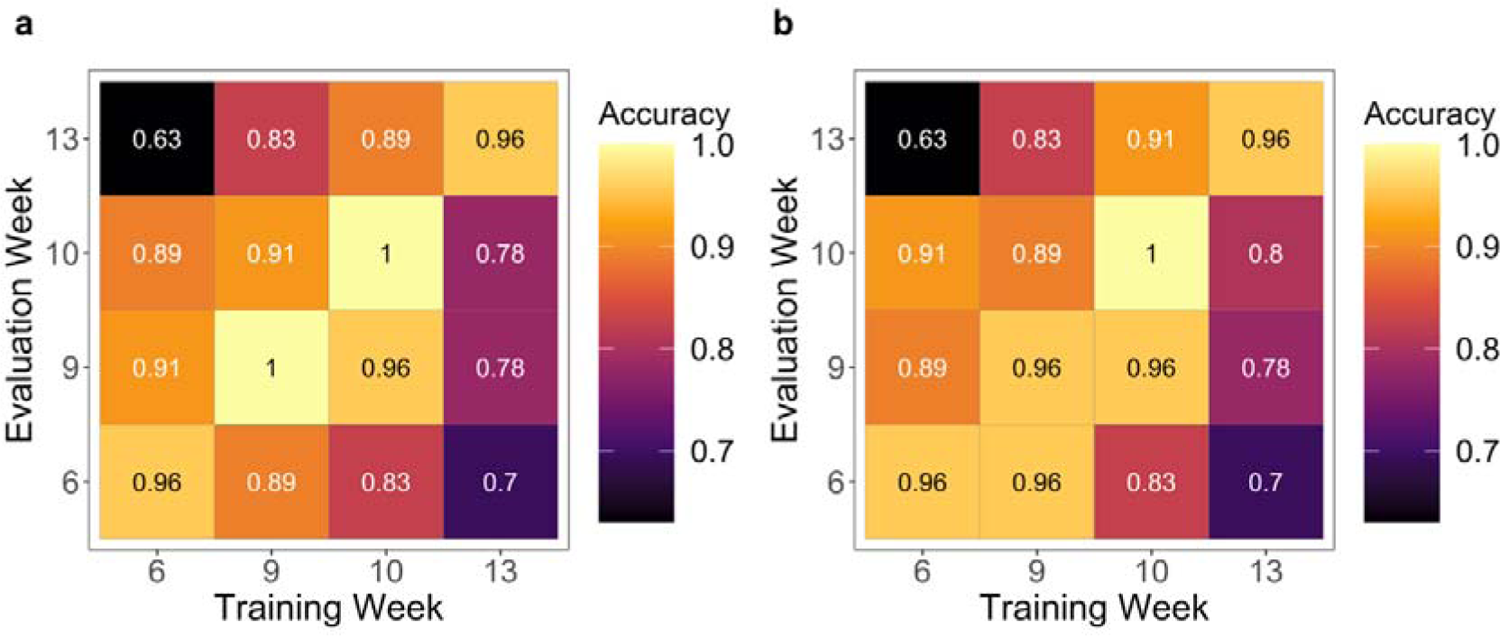
Nitrogen treatment levels predicted by support vector machines. (a) radial basis function kernels and (b) linear kernels in support vector machines, trained with W_SVM_ from the training week.

### Use of hyperspectral imaging as surrogates for physiological traits in rice

The ReliefF algorithm was used to derive W_PLSR_, *i.e.* wavelengths selected for partial least squares regression (PLSR, see Methods) to reduce the dimensionality of HSI data (**Figure 5a** and **Figure 6a**). Reflectance data of W_PLSR_ was the input of a single-response PLSR model. Among measured traits, models could be best developed for *N*, and *C:N,* followed by *SLA* (**Table 2**). The number of components used for *N* and *C:N* were six and seven, respectively (**Figure 5b** and **Figure 6b**). For the other traits, model metrics was not ideal with %RMSEP greater than 15% in validation datasets.

**Figure 5.**
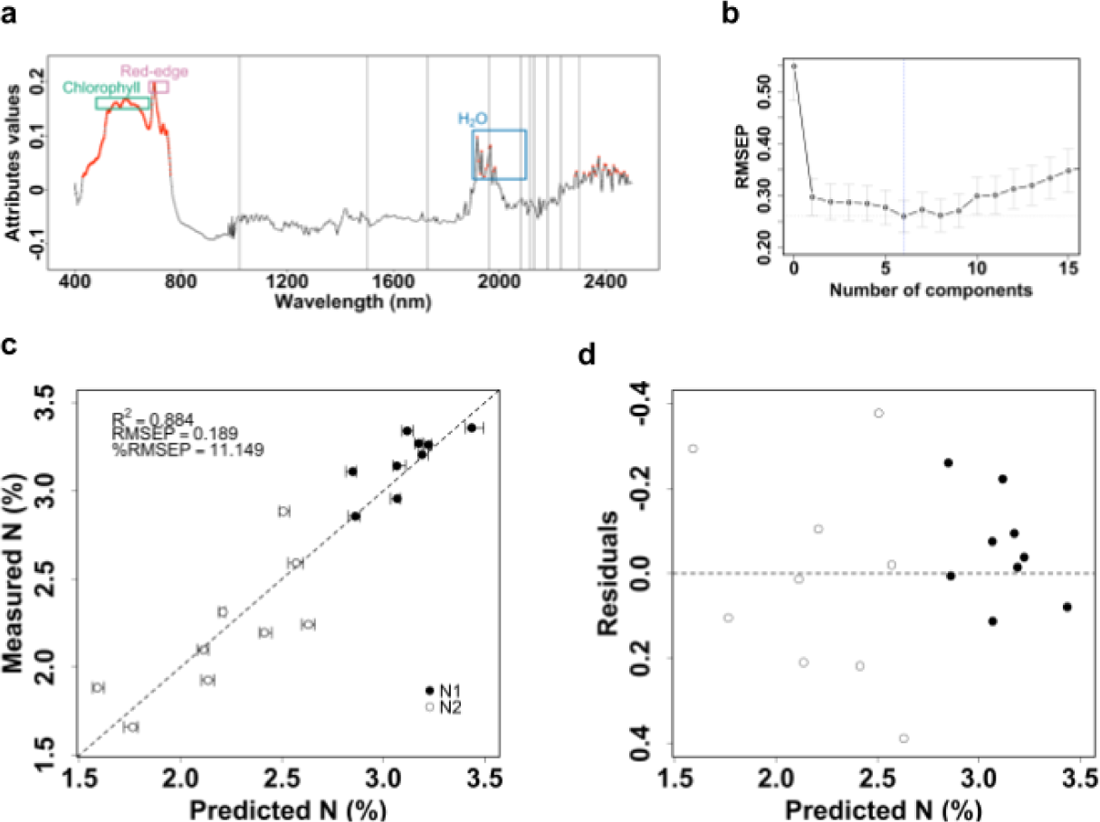
Predicting leaf-level nitrogen from HSI data. (a) Three hundred wavelengths (red) are important for leaf-level nitrogen (N, %) based on the attributes values calculated from ReliefF algorithm. The light gray vertical lines (1020, 1500, 1730, 1960, 2080, 2115, 2130, 2180, 2230 and 2300 nm) are wavelengths that are responsive to N-H stretch or proteins according to the research of Fourty *et al*. (1996) and Ecarnot *et al*. (2013), (b) model performance on validation dataset (n=18). The gray dashed line is 1:1 ratio line. Solid and empty circles are high nitrogen (N1) and low nitrogen (N2) treatments, respectively. The error bars are 95% confidence intervals derived from jackknife permutation and (c) residuals from the validation dataset.

**Figure 6.**
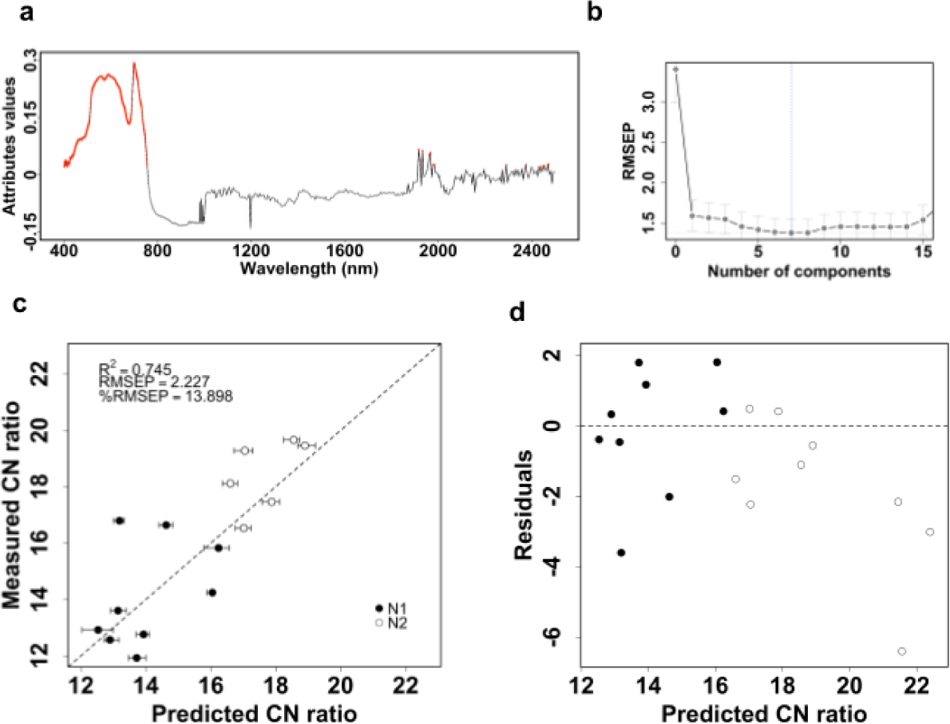
Predicting leaf-level carbon to nitrogen (C:N) ratio from HSI data. (a) Three hundred wavelengths (red) are important for CN ratio based on the attributes values calculated from ReliefF algorithm, (b) model performance on validation dataset (n=18). Solid and empty circles are high nitrogen (N1) and low nitrogen (N2) treatments, respectively. The gray dashed line is 1:1 ratio line. The error bars are 95% confidence intervals derived from jackknife permutation and (c) residuals from the validation dataset.

**Table 2.** Summary statistics for PLSR models. Model results of trait prediction using HSI data. The lowest RMSEP in the calibration dataset is used to choose the number of components in the model. NC, number of components, R^2^, coefficient of determination; RMSEP, root mean square of error in prediction; %RMSEP, normalized root mean square of error in prediction; Cal, calibration dataset; Val, validation dataset.

| Traits | N Cal | N Val | NC | $R^2$ Cal | RMSEP Cal | %RMSEP Cal | $R^2$ Val | RMSEP Val | %RMSEP Val |
| --- | --- | --- | --- | --- | --- | --- | --- | --- | --- |
| N | 70 | 18 | 6 | 0.77 | 0.26 | 11.09 | 0.88 | 0.19 | 11.15 |
| C:N ratio | 70 | 18 | 7 | 0.83 | 1.38 | 10.25 | 0.75 | 2.23 | 13.90 |
| SLA_C | 70 | 18 | 8 | 0.15 | 0.85 | 20.15 | 0.41 | 0.84 | 18.52 |
| SLA | 74 | 20 | 7 | 0.33 | 20.94 | 15.89 | 0.31 | 23.32 | 20.29 |
| mQY | 74 | 20 | 5 | 0.10 | 0.03 | 21.98 | 0.54 | 0.02 | 20.47 |
| E | 74 | 20 | 2 | -0.01 | 0.00 | 17.21 | -0.05 | 0.00 | 22.84 |
| $V_{cmax}$ | 70 | 19 | 1 | 0.00 | 19.80 | 19.47 | 0.25 | 13.88 | 24.41 |
| A | 75 | 19 | 1 | 0.04 | 0.51 | 13.67 | -0.44 | 0.56 | 24.69 |
| gsw | 74 | 20 | 2 | 0.00 | 0.13 | 18.02 | -0.13 | 0.11 | 24.70 |
| FB | 74 | 20 | 12 | 0.00 | 25.71 | 21.02 | -0.19 | 33.80 | 29.24 |
| $J_{max}$ | 70 | 19 | 7 | 0.06 | 52.10 | 16.24 | -1.02 | 51.04 | 35.74 |
| SLN | 70 | 18 | 3 | 0.14 | 0.02 | 13.05 | -0.73 | 0.01 | 36.46 |
| pQY | 74 | 20 | 3 | 0.313 | 0.021 | 15.539 | -0.995 | 0.026 | 45.90 |
| C | 70 | 18 | 0 |  |  |  |  |  |  |

PLSR using HSI data predicted *N* (**Figure 5**) and *C:N* (**Figure 6**) well with *R^2^* values of 0.88 and 0.75, respectively, and %RMSEP values of 11.15% and 13.90%, respectively (in validation datasets). It was not surprising to find that the wavelengths with high attributes values of *N* and *C:N* localized to similar regions (**Figure 5a** and **Figure 6a**). Specifically, the VIS region and the beginning of the NIR region were critical to predict both *N* and *C:N.* In addition, the attributes values had remarkable peaks in the Red Edge region of both traits. The performance metrics of *N* were higher in the validation dataset than the calibration dataset, which we hypothesized to be due to the stochastic nature of selecting samples for the validation and calibration datasets. This was confirmed by sampling multiple new validation and calibration datasets with different seed values.

## DISCUSSION

In this study, we leveraged genetic differentiation of two rice subpopulations along with two levels of nitrogen application to expand the range of observed phenotypic variation in a selected suite of plant growth and physiological traits. Our overarching goal was to assess the potential utility of hyperspectral reflectance information from automated imaging taken from a large-scale controlled-environment phenotyping facility to use as surrogates for laborious-to-measure plant traits. Due to the explicit use of both genetic and environmental variation in the experimental design phase, new insights into best practices for developing hyperspectral models for larger scale genetic studies may be gleaned from this work. The relatively long experimental period (13 weeks) additionally afforded a unique opportunity to ask about the persistence of HSI utility over the course of plant development.

Overall, we found that the traits, *N* (leaf-level nitrogen content) and *C:N* (leaf-level carbon to nitrogen ratio) could be predicted using HSI data, whereas for other traits, models either could not be developed or resulted in poor validation. Of note, *N* and *C:N* were traits that contributed most to the first principal component of the PCA using ground-truth data, which separated nitrogen treatment classes, and we speculate that this tight relationship contributed to the high performance of the PLSR models for these traits. Previous work has also demonstrated successful *N* prediction in crop species using HSI data (Vigneau et al., 2011; Din et al., 2017; Tan et al., 2018; Bruning et al., 2019; He et al., 2020; Meacham-Hensold et al., 2020; Vergara-Diaz et al., 2020; Yu et al., 2020; Baath et al., 2021; Lin et al., 2022). Under various growth conditions, changes in reflectances around 700 and 900 nm (Red Edge and NIR, respectively) have been identified as critical for trait prediction, and our findings were consistent with this (**Figures 3a, 5a and 6a**). In addition to the known utility of the Red Edge and NIR regions for *N* prediction, we found that wavelengths around 1800 and 2400 nm (SWIR region) contributed to the separation of nitrogen treatments (**Figure 3a** and **Table S4**), and according to attributes values, wavelengths around 580 nm were additionally important for predicting *N* and *C:N*. To our knowledge, only two recent studies have marked these regions as potentially associative to *N*. The work of Vergara-Diaz *et al*. (2020) identified wavelengths of 1680, 2050 and 2170 nm in the SWIR region to be responsive to nitrogen and protein content in durum wheat, and Yu *et al*. (2020) selected wavelengths at 500, 566, and 600 nm in the VIS region to model nitrogen differences in rice (Gengyou 653, a hybrid from *japonica* subpopulation). While there are dissimilarities in the specific wavelengths between these studies and ours, likely due to differences in sensors and/or study species/varieties, the previous results along with ours together provide evidence that the SWIR and VIS regions can contribute to the prediction of *N*.

Grouping the ground-truth measurements into trait types (Group 1: biomass constituents [*N, C,* and *C:N*]; Group 2: weights and ratios of weights and area [*FB, SLN, SLA* and *SLA_C*]; Group 3: fluxes [*A, E, gsw*]; and Group 4: photosynthetic capacity [*J_max_, V_cmax_, pQY, mQY*]), we note that both predicted traits, *N* and *C:N,* are biomass constituents. However, HSI data were unable to predict *C*, also part of Group 1. Insufficient variation in *C* in our plants may have led to the poor model performance. Indeed, the coefficients of variation for *C* and *N* were 0.0474 and 0.2020, respectively, indicating that the variation of *C* was much lower than that of *N*. For Group 2 traits, *FB* was unable to be predicted using our HSI data. This sits in contrast to previous research that has leveraged HSI to predict biomass (Cho et al., 2007). One critical distinction is that the previous study used fresh weight as the metric for biomass. As water constitutes 90% of the mass in the growing tissue of a plant cell (Chavarria and dos Santos, 2012) and hyperspectral reflectance is sensitive to water (Gewali et al., 2019), it is likely that dry biomass is hard to predict from HSI data collected during the period plants are alive. Instead, dry weight may be better predicted using an alternative high-throughput method such as RBG imaging that can readily capture morphological traits (e.g., top surface area) that are highly correlated with biomass (Pandey et al., 2017). For the other traits in Group 2 as well as traits in Groups 3-4, we surmise that one general underlying reason for the inability to build satisfactory models is due to the differences in spatial scale between imaging data (taken at canopy-level angles) and ground-truth measurements (taken at leaf-level of the youngest fully expanded leaf). For these particular traits, heterogeneity among individual leaves is likely present, which would not have been captured by our ground-truth measurements. Previous research successfully predicted *SLA* from HSI data by utilizing a hand-held spectroradiometer on the same individual leaves used for ground-truth measurements (Cotrozzi et al., 2020). In addition, as the rice plants grew, the architecture of the plants became increasingly complex, which may mask potential physiological information in the HSI data due to complicated interaction between plant tissue and light (Al Makdessi et al., 2019; Asaari et al., 2019; Mishra et al., 2020; Mertens et al., 2021). To improve model performance, future work using canopy-level HSI systems should strive to capture some intra-canopy variation in ground-truth trait data. Similarly, temporal lags between ground-truth measurements and imaging events should be minimized, especially for Group 3 traits that represent fluxes, which are very sensitive to environmental differences. In our particular setup at the AAPF, plants must travel on conveyor belts out of the growth chamber environment to the imaging booth (**Figure S7**). This takes approximately four minutes, during which temperature dropped to around 21 °C. Ge *et al*. (2016) and Yang *et al*. (2020) also pointed out that photosynthesis processes were likely to be different between the growing environments of plants and imaging booths. Ge *et al*. (2016) suggested that a protocol to minimize the time it takes for plants to travel from growth chambers to imaging systems should be implemented if the researchers want to gain understanding of photosynthetic traits from the imaging data. Yang *et al*. (2020) suggested that using a sensors-to-plants approach could also alleviate the effects of temporal decoupling.

One question we were keen to address here was whether HSI signatures differed in accordance with subpopulation identity. We determined that, despite clear subpopulation differentiation in trait data, there were no background genetic signatures in HSI information for rice, based on PCA and model predictions *of N* and *C:N.* This suggests several implications for moving forward in conducting larger-scale experiments leveraging HSI-predicted physiological traits in diverse rice: (1) subpopulation genetic background does not need to be controlled for in prediction models; and (2) calibration models can be developed from a combination of genetic materials, similar to NIR models for stem non-structural carbohydrate traits in rice (Wang et al., 2016, 2017) (conditional that there is adequate phenotypic variation). Contrary to our results, He *et al*. (2020) suggested that *indica* and *japonica* subpopulations required different models to estimate *N*. The main difference could be that in their study, HSI data was taken at the top-view whereas our analyzed HSI data was side-view data. This may suggest viewing angles of the cameras affect whether model calibration needs to consider genotypic variation.

Automated imaging systems, such as Purdue’s AAPF, readily support high frequency hyperspectral reflectance measurements, as imaging events can be programmed prior to the start of the experiment and require limited human intervention (**Figure S7**). In our study, imaging events occurred two to three times *per* week throughout the entire experimental period. The frequency of imaging allowed us to average across several days to obtain pot-level spectral information to be used for downstream analysis and modeling; this attenuates potentially noisy individual spectra and helps detect the overall HSI signature of each plant during timepoints of interest (**Table S2**). Having automated imaging occur from Week 6 through Week 13 also enabled us to ask about the consistency of HSI information over developmental time. To test this, we employed SVM to see if HSI could classify nitrogen treatments forwards and backwards in time (*i.e*., using one week to predict another week). Classification using HSI data performed very well overall (**Figure 4**), however, it was interesting to find that the latest timepoint, Week 13, had the worst performance out of all weeks, whether it served as the Training or the Evaluation Week. This timepoint occurred at the very end of the experimental period, and we speculate that water may have played an interactive role with nitrogen treatment to modify HSI signatures. We found that higher levels of nitrogen led to greater *SLA* in our experiment (**Table S3**), meaning greater transpiring leaf area *per* unit dry biomass; this translates to an increased demand for water. Watering regime, which was managed on a target weight basis using the facility’s automated system, did not differ between the two nitrogen treatments; therefore, it is plausible that the high nitrogen treatment would have experienced some level of water stress by the end of the experiment since it had greater *SLA*. This explanation is supported by two additional observations: (1) by Week 13, key wavelengths derived from PCA showed a shift from 2400 nm to 1800 nm, adjacent to the region known to be responsive to water (**Figure 3a**) and (2) we noticed leaf-rolling, a sign of water stress (O’Toole and Cruz, 1980), in some plants during midday in the later part of the experimental period.

One challenge to fully leveraging hyperspectral information for plant science research lies in being able to select useful information from data characterized by an abundance of highly collinear variables (Baath et al., 2021). Reducing the dimensionality of HSI processed output is therefore a key step towards successfully modeling plant traits (Gewali et al., 2019). For this purpose, various ML algorithms have been proposed, as they are flexible and adaptive to the data collected (Verrelst et al., 2015; Gewali et al., 2019; Mohd Asaari et al., 2022). Our current study utilized PCA, SVM, ReliefF algorithm and PLSR methods. We demonstrated that PCA and ReliefF were useful methods to reduce the dimensionality of HSI data.

The promise of leveraging high-throughput and non-destructive HSI data as surrogates for plant physiological traits has motivated a wealth of research to test the applicability of HSI across a variety of crops and treatments (Fiorani and Schurr, 2013). However, variation in these data from automated plant phenotyping facilities is a function of multiple factors such as changes in plant structure through time, scaling, equipment (e.g., cameras) or the particular setup of the facility, which impacts the timing of imaging as plants move from their growing environment to imaging booths. For these reasons, prediction models are generally non-transferrable across species, experiments, or facilities (Cho et al., 2007; Verrelst et al., 2015). Flexible approaches to designing training sets with associated ground-truth data particular to the experiment at hand are therefore needed. With appropriate training sets, machine learning algorithms can then be leveraged to filter high-dimensional information from HSI data and to generate relationships between input and output. Importantly, researchers need to examine how HSI and ground-truth data are collected, with spatial and temporal considerations in mind. Finally, plant scientists may want to collaborate closely with imaging facility engineers and technicians over the lifetime of an experiment, from experimental design to data analysis. Improvements made in this direction are critical if HSI technologies are to lead to significant biological insights in the plant sciences.

## SUPPLEMENTARY DATA

Table S1 Germplasm information for the rice panel

Table S2 Timing of physiological trait measurements and imaging dates

Table S3 Summary statistics of physiological traits

Table S4 Wavelengths selected for support vector machines from Week 6 to Week 13

Table S5 Summary of weekly Vegetation Indices

Figure S1 Geographical distribution of the selected rice accessions in the study

Figure S2 Growth conditions inside the Ag Alumni Phenotyping Facility

Figure S3 Histograms overlaid with parameterized normal density curves of the measured physiological traits

Figure S4 Comparison of measured physiological traits

Figure S5 Pairwise relationship among the measured physiological traits

Figure S6 Wavelength profiles for each rice plant across days

Figure S7 Rice plants on the inspection belt at the Ag Alumni Phenotyping Facility at Purdue University (West Lafayette, Indiana, U.S.A.)

## ABBREVIATIONS

A: (pre-dawn leaf-level net assimilation)

C: (carbon)

C:N: (leaf-level carbon to nitrogen ratio)

E: (pre-dawn leaf-level transpiration)

(FB): final biomass

gsw: (pre-dawn leaf-level stomatal conductance to water)

HSI: (hyperspectral images)

J_max_: (light-saturated potential rate of electron transport)

log(E): (log transformed pre-dawn leaf-level transpiration)

log(gsw): (log transformed pre-dawn leaf-level stomatal conductance to water)

mQY: (midday quantum yield)

%RMSEP: (normalized root mean square of error in predictions)

NIR: (near infrared)

N: (nitrogen)

(PLSR): partial least squares regression

pQY: (pre-dawn quantum yield)

RMSEP: (root mean square error in predictions)

RBF: (radial basis function)

RGB: imaging (Red-Green-Blue imaging)

*R^2^*: (coefficient of determination)

SLA: (specific leaf area with respect to dry weight)

SLA_C: (specific leaf area with respect to carbon)

SLN: (specific leaf nitrogen)

SVM: (support vector machine)

SWIR: (shortwave infrared)

V_cmax_: (the maximum carboxylation rate limited by Rubisco)

VIS: (visible)

Wplsr: (wavelengths selected for PLSR)

Wsvm: (wavelengths selected for model input of SVM)

## ACKNOWLEDGEMENTS

The authors thank Makala Hammons and Natalie Roth for help with plant measurements, Yujie Chen for statistical consulting, and John Couture and Yiwei Jiang for feedback on analyses.

## AUTHOR CONTRIBUTIONS

AS, CRG, CH, YY and DRW conceptualized the study. RKI, CH, and DRW collected data. AS and TT analyzed data. TT and DRW wrote the manuscript and all authors contributed feedback and/or edits.

## DATA AVAILABILITY

The datasets collected and analyzed for this study can be found in the Purdue University Research Repository [https://purr.purdue.edu/publications/4079/1].

